# Loss of *Tsc1* in cerebellar Purkinje cells induces transcriptional and translation changes in FMRP target transcripts

**DOI:** 10.1101/2021.03.03.433717

**Authors:** Jasbir Dalal, Kellen D. Winden, Catherine L. Salussolia, Maria Sundberg, Achint Singh, Pingzhu Zhou, William T Pu, Meghan T. Miller, Mustafa Sahin

**Author notes:** Corresponding author: Mustafa Sahin. MTM is now an employee of Skyhawk Therapeutics. These authors contributed equally to this manuscript (Jasbir Dalal, Kellen D. Winden).

## Abstract

Tuberous sclerosis complex (TSC) is a genetic disorder that is associated with multiple neurological manifestations. Previously, we demonstrated that *Tsc1* loss in cerebellar Purkinje cells (PCs) can cause altered social behavior in mice. Here, we performed detailed transcriptional and translational analyses of *Tsc1*-deficient PCs to understand the molecular alterations in these cells. We found that target transcripts of the Fragile X Mental Retardation Protein (FMRP) are reduced in mutant PCs with evidence of increased degradation. Surprisingly, we observed unchanged ribosomal binding for many of these genes using Translating Ribosome Affinity Purification (TRAP). Finally, we found that the FMRP target, SHANK2, was reduced in PC synapses, suggesting that compensatory increases in ribosomal binding efficiency may be unable to overcome reduced transcript levels. These data further implicate dysfunction of FMRP and its targets in TSC and suggest that treatments aimed at restoring the function of these pathways may be beneficial.

## Introduction

Tuberous sclerosis complex (TSC) is a neurocutaneous disorder caused by germline heterozygous loss-of-function mutations in *TSC1* or *TSC2*. Among the neurological manifestations, approximately 50% of patients with TSC are diagnosed with autism spectrum disorder (ASD) (Jeste et al., 2008; Jeste et al., 2016; Capal et al., 2017). Disrupted development of the cerebellum is among a number of pathogenic processes that have been associated with ASD (Fatemi et al., 2012; Becker and Stoodley, 2013; Rogers et al., 2013), and cerebellar pathology has been identified in patients with TSC (Weber et al., 2000; Eluvathingal et al., 2006; Vaughn et al., 2013; Weisenfeld et al., 2013). We have previously demonstrated that loss of either one or both alleles of *Tsc1* in cerebellar Purkinje cells (PCs) results in abnormal social behavior (Tsai et al., 2012; Tsai et al., 2018). Therefore, we evaluated the molecular targets that are dysregulated in TSC1-deficient PCs to understand the mechanism by which loss of TSC1 leads to PC dysfunction and abnormal social behavior.

TSC1 and TSC2 form a protein complex (TSC1/2) that negatively regulates the mechanistic target of rapamycin (mTOR) pathway by inactivating RHEB. Both TSC1 and TSC2 are necessary for a functional TSC1/2 complex (Hoogeveen-Westerveld et al., 2011), and loss of either gene leads to the clinical manifestations of TSC that are indistinguishable from one another. Thus, loss of either TSC1 or TSC2 leads to unchecked activation of mTOR. mTOR is a kinase that regulates several cellular processes, including proliferation, protein synthesis, and autophagy (Lipton and Sahin, 2014; Liu and Sabatini, 2020). The phosphorylation targets of mTOR depend on its surrounding adapter proteins, which define either the mTOR complex 1 (mTORC1) or mTOR complex 2 (mTORC2). mTOR signaling from either complex may affect transcriptional regulation of several genes and pathways, and we previously observed that loss of TSC2 in cortical neurons caused upregulation of the transcription factor ATF3 (Nie et al., 2015). However, mTORC1 also facilitates translation initiation and elongation through phosphorylation of 4EBP1/2 and ribosomal protein S6 (Saxton and Sabatini, 2017). Therefore, loss of TSC1/2 can cause complex changes in protein expression through modulation of transcription and translation in neurons.

In this study, we sought to characterize the effects of loss of *Tsc1* on PCs at both the level of transcription and translation. We performed RNA sequencing of total RNA from sorted PCs using FACS, and we used Translating Ribosome Affinity Purification (TRAP) to determine ribosomal binding of transcripts in PCs. By comparing these different levels of RNA regulation, we observed that many transcripts that bind to FMRP maintain ribosomal binding despite reduced levels of transcripts overall. For one gene, *Shank2*, we found that the result of these opposing effects was reduced levels at PC synapses. These data highlight a complex interplay between the effects of mTOR activation of the regulation of transcript levels and ribosomal binding and provide further insight into the dysfunction of PCs in TSC.

## Results

### Loss of *Tsc1* in PCs leads to down-regulation of gene expression

To assess the effect of loss of *Tsc1* on the transcriptome of PCs, we designed an experimental paradigm that allowed us to isolate PCs and measure their gene expression. We utilized our previously characterized animal model where *Tsc1* harbors loxP sites flanking exons 17 and 18, which leads to a null allele after Cre-mediated recombination (Kwiatkowski et al., 2002; Meikle et al., 2007). In this model, Cre is driven by the L7 (or PCP2) promoter that leads to expression of Cre specifically within PCs (Tsai et al., 2012). We then crossed these animals with an animal that expresses GFP under the same L7 promoter, leading to expression of GFP specifically within PCs (Tomomura et al., 2001). We performed immunostaining of control (*Tsc1*^w/w^ L7GFP or WW) and mutant (*Tsc1*^c/c^ L7Cre;L7GFP or CC) animals. We observed that PCs were specifically labeled with GFP and that loss of *Tsc1* led to increased soma size and phosphorylation of S6, consistent with mTORC1 activation (Figure 1a-b).

**Figure 1:**
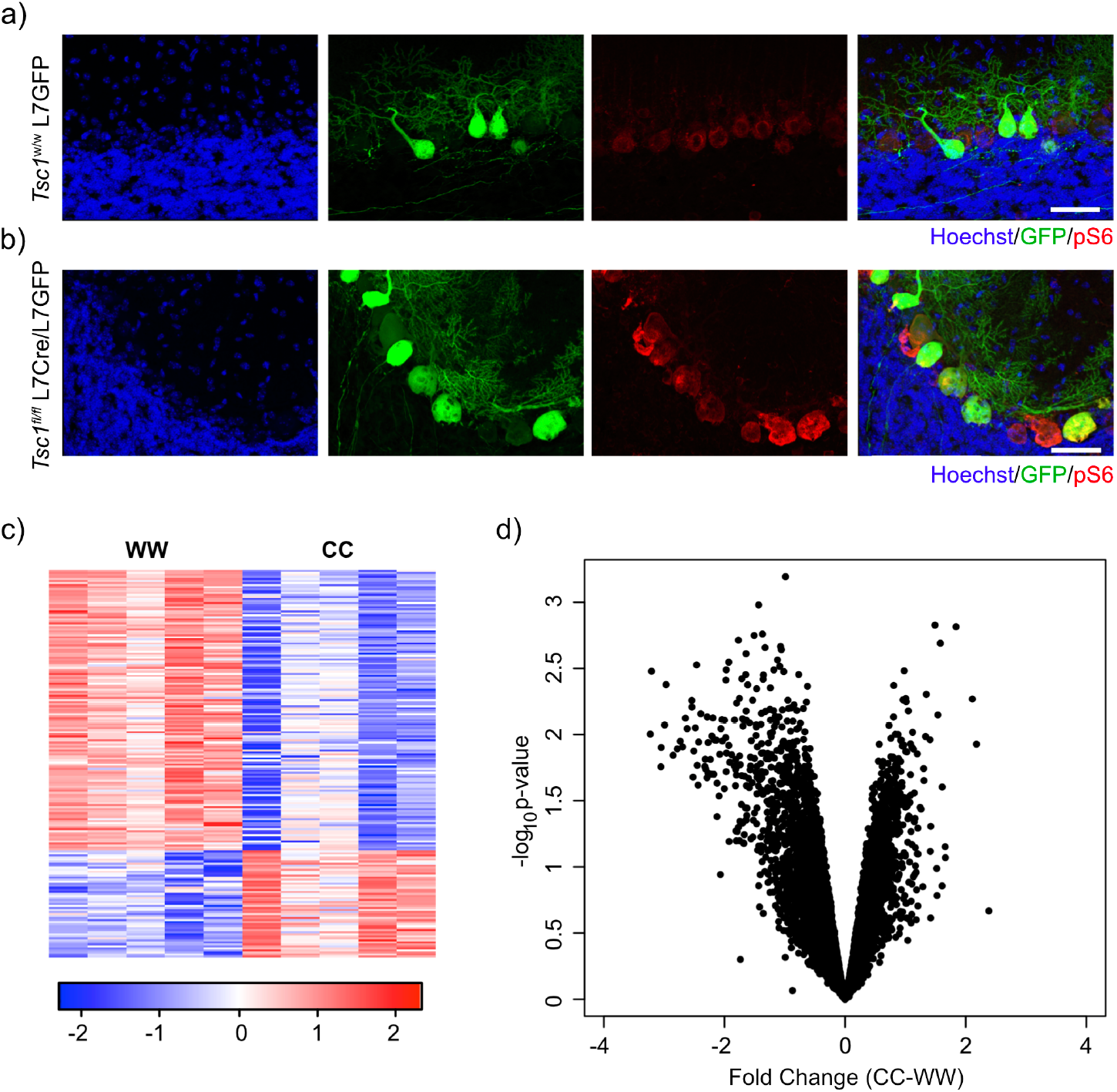
Labeling and sorting *Tsc1*^c/c^ PCs shows predominance of down-regulation gene expression. a) Immunocytochemical characterization of wildtype control L7-GFP^+^ mouse PCs on P21 mouse cerebellum. Cell nuclei were stained with Hoechst. GFP^+^ PCs had dim expression of phosphorylated S6 (red). Scale bar 50 mM. b) Immunocytochemical characterization of Tsc1^f/f^Cre^+^L7-GFP ^+^ mouse PCs on P21 mouse cerebellum. Cell nuclei were stained with Hoechst. GFP^+^ PCs were strongly positive for pS6 (red). Scale bar 50 mM. c) Heatmap of differentially expressed genes in mutant (*Tsc1*^c/c^;N=4) vs control (*Tsc1*^w/w^;N=4) PCs (FC > 2 and p-value < 0.01; n=679 genes). Each row of the heatmap represents the scaled expression of one gene, where red corresponds to higher relative expression and blue corresponds to lower relative expression. d) This volcano plot shows the relationship between fold change, calculated *Tsc1*^c/c^ – *Tsc1*^w/w^, and p-value across all genes. A majority of genes in this dataset demonstrate down-regulation in the mutant PCs compared to control.

We then dissociated the cerebellum of control (N=4) and mutant (N=4) animals into single cell suspensions and performed fluorescence activated cell sorting (FACS) to specifically isolate GFP-positive PCs. After removal of debris, doublets, and dead cells, we found that GFP-positive PCs comprised approximately 0.5% of total events (Supplemental figure 1). We then isolated total RNA from these samples and performed RNA sequencing. Differential expression analysis between control and mutant animals identified 192 differentially expressed genes (FC > 2 and p-value < 0.01; Supplemental table 1). Interestingly, most of the differentially expressed genes in this dataset were down regulated (72%) (Figure 1c). Indeed, we found that there was a bias towards down-regulation in the overall gene expression, even among genes that did not show a significant change in gene expression (Figure 1d). This effect was more evident in genes with higher expression, and there was a statistically significant negative correlation between fold change and expression (r=-0.14, p<2.2e-16, Pearson correlation). These data demonstrate that loss of *Tsc1* results in a bias towards widespread transcriptional down-regulation.

### Increased degradation of down-regulated transcripts in *Tsc1*^c/c^ PCs

We hypothesized that the bias towards reduced levels of a majority of transcripts may reflect a change in RNA stability caused by loss of *Tsc1*. Therefore, we asked whether there was a systematic alteration in the coverage of transcripts from the RNA sequencing data to understand whether RNA degradation pathways might be altered in mutant PCs. We used the aligned RNA sequencing data to calculate the coverage of each nucleotide across each transcript, and we focused on transcripts that showed an average coverage of at least one read per nucleotide. We then divided each transcript into 100 equally sized intervals, and we calculated the fraction of reads within each interval compared to the total number of reads across the transcript. When we compared the median coverage across all transcripts between mutant and control PCs, we observed that mutant PCs showed reduced coverage at the 3’ end of the transcript compared to control PCs (Figure 2a). These data show that there is a bias towards reduced coverage in the 3’ region of transcripts in mutant PCs, suggesting that 3’->5’ RNA degradation pathways may be increased in mutant PCs.

**Figure 2:**
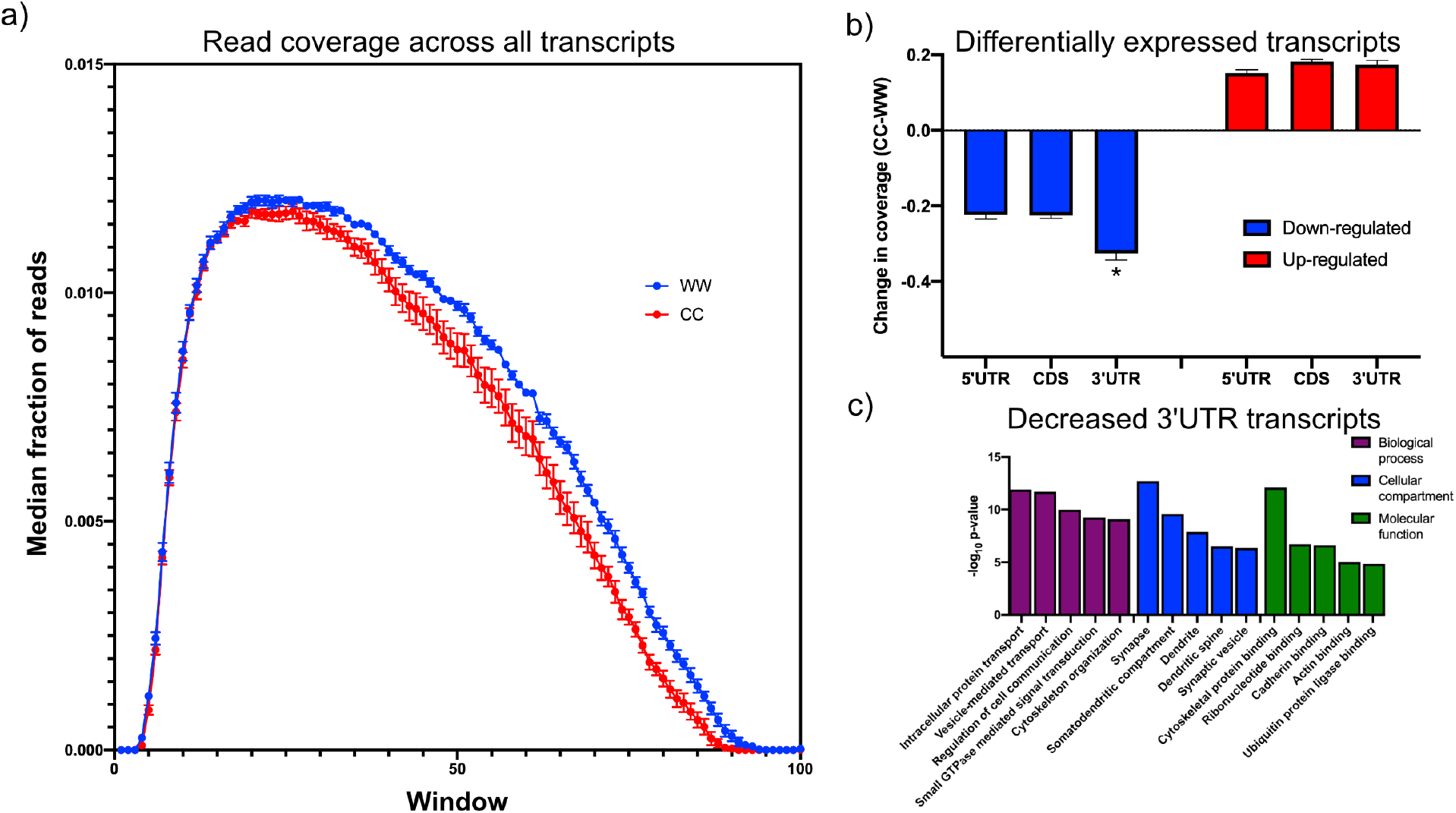
Down-regulated transcripts in Tsc1-KO PCs display enhanced reduction in the 3’UTR and are enriched in synaptic genes. a) This plot shows the median coverage of reads across all down-regulated transcripts. Each transcript was divided into 100 equally sized intervals and coverage was calculated across each of these intervals. The error bars show the SEM between five animals of each genotype, where mutant PCs are shown in red and control PCs are shown in blue. There is a bias for fewer reads across the 3’ regions of transcripts in mutant PCs. b) This bar plot shows the read coverage across different segments of each transcript for down-regulated (blue) and up-regulated (red) transcripts. Read coverage was calculated separately for the 5’ UTR, coding sequence, and 3’ UTR, and the error bars represent SEM. The 3’ UTR of down-regulated genes in mutant PCs demonstrate significantly reduced coverage compared to the 5’ UTR and coding sequence (*p<0.001, ANOVA with Tukey post-hoc test). c) This bar plot shows the enrichment of functional gene ontology categories among the down-regulated transcripts with enhanced reduction in the 3’ UTR. Several synapse-related categories are among the most enriched categories.

To further understand the dependence between overall transcript expression and coverage across the transcript, we identified all significantly up- and down-regulated transcripts and analyzed the difference in coverage of these genes between mutant and control across the 5’ UTR, the coding sequence, and the 3’ UTR to determine whether changes in one of these regions was responsible for driving the change in expression of the transcript. The 3’ UTR showed significantly reduced coverage in the mutant compared to the control PCs among the down regulated genes (Figure 2b). However, there was no significant difference between the regions of the up-regulated genes. These data suggest that there may be distinct mechanisms driving up-and down-regulation of genes within TSC1-deficient PCs.

We then searched through the down-regulated transcripts for those that showed greater changes in the 3’ UTR than the overall transcript expression, identifying 978 transcripts (Supplemental table 2). We reasoned that this may be a core group of transcripts that are strongly affected by increased RNA degradation in mutant PCs. To understand what functions are associated with this group, we performed gene ontology analysis and observed that these transcripts were strongly enriched in categories such as synapse, protein transport, and cytoskeletal binding (Figure 2c). In a prior study, we had observed that TSC2-deficient PCs that were derived from human induced pluripotent stem cells showed down-regulation of FMRP targets (Sundberg et al., 2018). Therefore, we examined this group of down-regulated transcripts and found a highly significant enrichment of FMRP target genes (p=2.59e-55)(Darnell et al., 2011). Taken together, these data demonstrate that down-regulated genes with alterations in the 3’ UTR are FMRP targets with associated synaptic functions, suggesting that perturbation of RNA decay pathways in mutant PCs may substantially affect PC physiology.

### TRAP of *Tsc1*^c/c^ PCs does not reflect decreased total transcript levels

mTOR also exerts strong effects at the level of translation, and therefore, we decided to evaluate ribosomal binding of transcripts within PCs to determine whether changes in translation mechanisms could compensate for or exacerbate the alterations that we had observed in the transcriptome. The TRAP paradigm utilizes a labeled ribosomal subunit (RPL10A-EGFP) that can be expressed in a subset of cells to specifically profile ribosomal binding of transcripts (Heiman et al., 2014). For these studies, we used animals that have a floxed-STOP RPL10A-EGFP in the constitutively active Rosa26 locus, which leads to EGFP-labeled ribosomes after Cre-mediated recombination (Figure 3a). We crossed these animals to the *Tsc1*/L7Cre animals described above. We then performed immunostaining of these animals, and we found that GFP was expressed specifically within calbindin-positive neurons, which is a known marker of PCs (Figure 3b).

**Figure 3:**
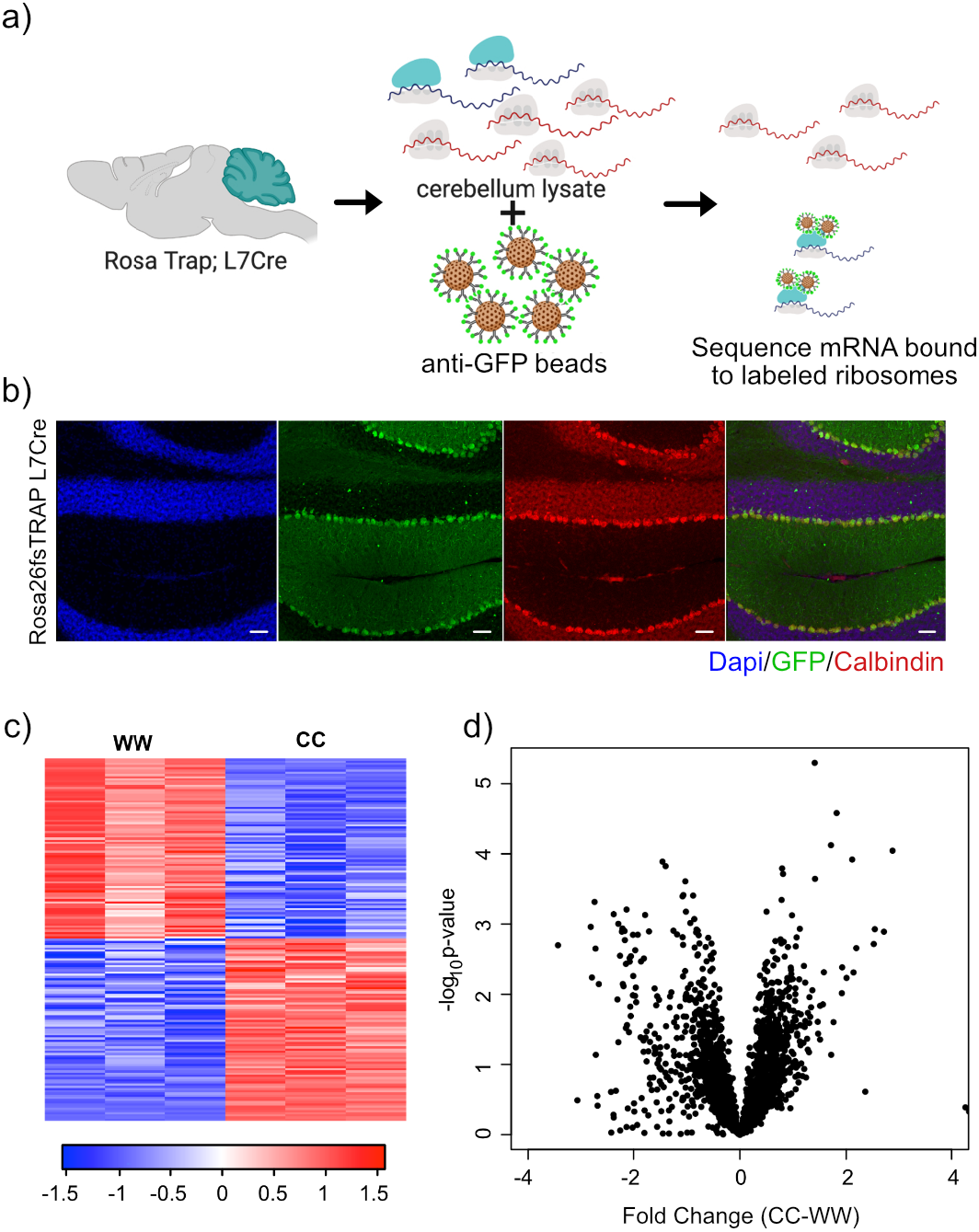
Expression of GFP-tagged ribosomes and ribosomal binding of transcripts in PCs. a) Schematic of TRAP protocol for identifying ribosomally-bound transcripts specifically within PCs. b) Anti-GFP immunofluorescence of Pcp2-eGFP/Rpl10a shows labeling of the PCs of the cerebellum. Sagittal sections from P56 Rosa-Trap animals expressing Cre under Pcp2 promoter were stained with GFP (green), calbindin (red), and DAPI (blue). Scale bars, 50um. c) Heatmap of TRAP levels of genes that were previously identified to be down-regulated at the transcript level. Each row of the heatmap represents the scaled expression of one transcript, where red corresponds to higher relative expression and blue corresponds to lower relative expression. d) This histogram shows the fold changes of the TRAP values for the previously identified down-regulated transcripts between mutant and control PCs. Despite being down-regulated at the total mRNA level, these transcripts generally show maintained ribosomal binding.

We performed RNA sequencing on TRAP samples from mutant (N=3) and control PCs (N=3). We then focused on genes that showed significant changes in ribosomal binding between mutant and control PCs. This analysis identified 165 differentially bound genes (FC > 2 and p-value < 0.01; Supplemental table 3). Interestingly, nearly equal numbers of genes showed increased and decreased ribosomal binding compared to control (Figure 3c). Consistent with these data, we found that there was no overall bias towards down-regulation of ribosome bound mRNAs (Figure 3d), in contrast to the total RNA data obtained from sorted PCs.

### Translation efficiency is increased in *Tsc1*^c/c^ PCs

mTOR is an important regulator of multiple aspects of translation, including initiation, elongation, and ribogenesis. Therefore, we examined ribosomal loading by calculating the translation efficiency (TE) for each gene, which is the difference between the ribosomal bound level of a gene and its total abundance in the cell (Thoreen et al., 2012). We then calculated the change in TE (ΔTE) between mutant and control animals for each gene. To estimate the variability of TE and determine whether changes in TE were statistically significant, we calculated ΔTE on subsets of the data and calculating the Z-score of each distribution. We found 1,808 genes that showed a ΔTE Z-score greater than 4, which corresponded to an uncorrected p-value < 3.2e-5 (Supplemental table 4). A majority of these genes (1,131) showed an increase in ΔTE in mutant PCs compared to control (Figure 4a). These data demonstrate that activation of mTOR through loss of TSC in PCs leads to increased TE, consistent with known roles of the mTOR pathway in facilitating protein synthesis.

**Figure 4:**
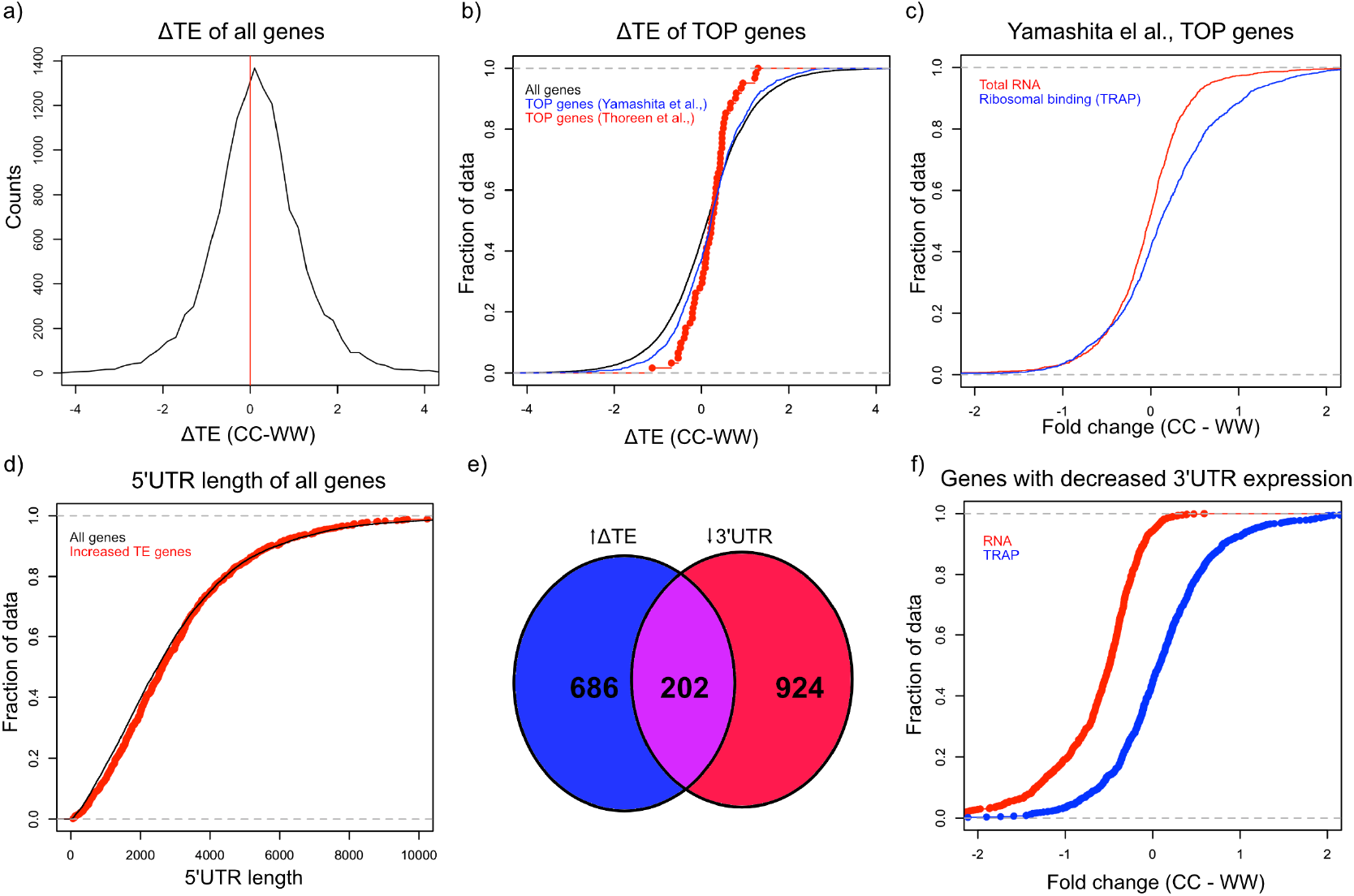
Alterations in TE in *Tsc1*^c/c^ PCs. a) This histogram shows the ΔTE between CC and WW PCs, and the vertical red line in the middle identifies the point where ΔTE=0. A majority of the genes are situated to the right of the red line demonstrating increased ΔTE in CC PCs. These empiric cumulative distribution function (ECDF) plots show the distribution for all genes (black), known TOP genes from Thoreen et al., (red), and predicted TOP genes from Yamashita et al., (blue). There is a slight shift towards higher ΔTE for Yamashita et al., predicted TOP genes compared to all genes (p=0.004, T-test). c) ECDF plot of fold changes for the total RNA and TRAP data for the predicted TOP genes from Yamashita et al., demonstrating that these genes have a slightly higher ribosomal binding in CC PCs without a change in the total RNA levels. d) ECDF plot of the 5’ UTR length for all genes (black) and genes with significantly increased ΔTE (red). There is no difference between these two groups of genes (p>0.05, Mann-Whitney U-Test). e) Venn diagram showing the highly significant overlap between genes with significantly increased ΔTE and down-regulated transcripts with reduced coverage in the 3’ UTR. f) ECDF plot of fold changes for the total RNA and TRAP data for down-regulated transcripts with reduced 3’ UTR coverage. These data show that the increased ΔTE of these genes is due to reduced total RNA levels with maintained ribosomal binding levels.

Known downstream targets of mTOR impact several aspects of protein synthesis in general, but studies have also found that mTOR can influence translation of specific transcripts. One specific motif that has been shown to increase translation in an mTOR related manner is the 5’ terminal oligopyrimidine (5’ TOP) motif, which is a pyrimidine rich motif near the transcription start site that is thought to facilitate initiation (Thoreen et al., 2012). Therefore, we searched for 5’ TOP motifs in the group of genes in mutant PCs that show increased TE. Although known TOP genes from one study (Thoreen et al., 2012) did not demonstrate a significant increase in TE, we found that all genes with a predicted TOP motif (Yamashita et al., 2008) displayed a significantly greater ΔTE compared to all other genes (p=0.004, T-test; Figure 4b). Consistent with this increase in TE, we found that these predicted TOP genes showed an overall positive fold change in ribosomal binding without a clear shift in total RNA abundance (Figure 4c). Other studies have found that length and/or complexity in the 5’ UTR region was an important feature of mTOR-regulated genes (Gandin et al., 2016). However, we found that there was no increase in the length of the 5’ UTR of transcripts with increased ΔTE (Figure 4d). We then examined the down-regulated genes with exaggerated reduction of the 3’ UTR identified above, and we found a remarkably significant overlap between these genes and those with significantly increased ΔTE (Figure 4e; p=2.08e-133; Hypergeometric probability). Interestingly, as opposed to the predicted TOP genes, these genes display down-regulation at the total transcript level and unchanged ribosomal binding, leading to their increased TE (Figure 4f). These data indicate that genes with enhanced down-regulation of the 3’ UTR, which are enriched in FMRP targets, show increased relative ribosomal binding with loss of TSC1.

### Increased TE of FMRP targets in *Tsc1*^c/c^ PCs

Given our observations that FMRP targets tended to be down-regulated at the transcript level but maintain their ribosomal binding, we decided to more carefully examine FMRP targets in TSC1-deficient PCs. We identified a group of 150 FMRP target genes based on known FMRP binding in prior studies (Darnell et al., 2011) and decreased expression in *Tsc1*^c/c^ L7Cre PCs, and we re-assessed their transcript level expression and RNA binding using amplicon-based quantification. We created amplicon-based libraries from total RNA from FACS PCs and TRAP RNA so that these data would be directly comparable without technical confounding due to different library preparation methods. In addition, the use of amplicons allowed interrogation of the same region of the transcript in both total and TRAP RNA. Consistent with our prior data, we confirmed that most of the FMRP targets were down-regulated in mutant PCs compared to control PCs (Figure 5a). However, these same genes showed no significant change in overall distribution in their ribosomal binding (Figure 5b). Therefore, these data confirm our observation of differential effects of loss of *Tsc1* on FMRP targets at the transcript and ribosome bound levels.

**Figure 5:**
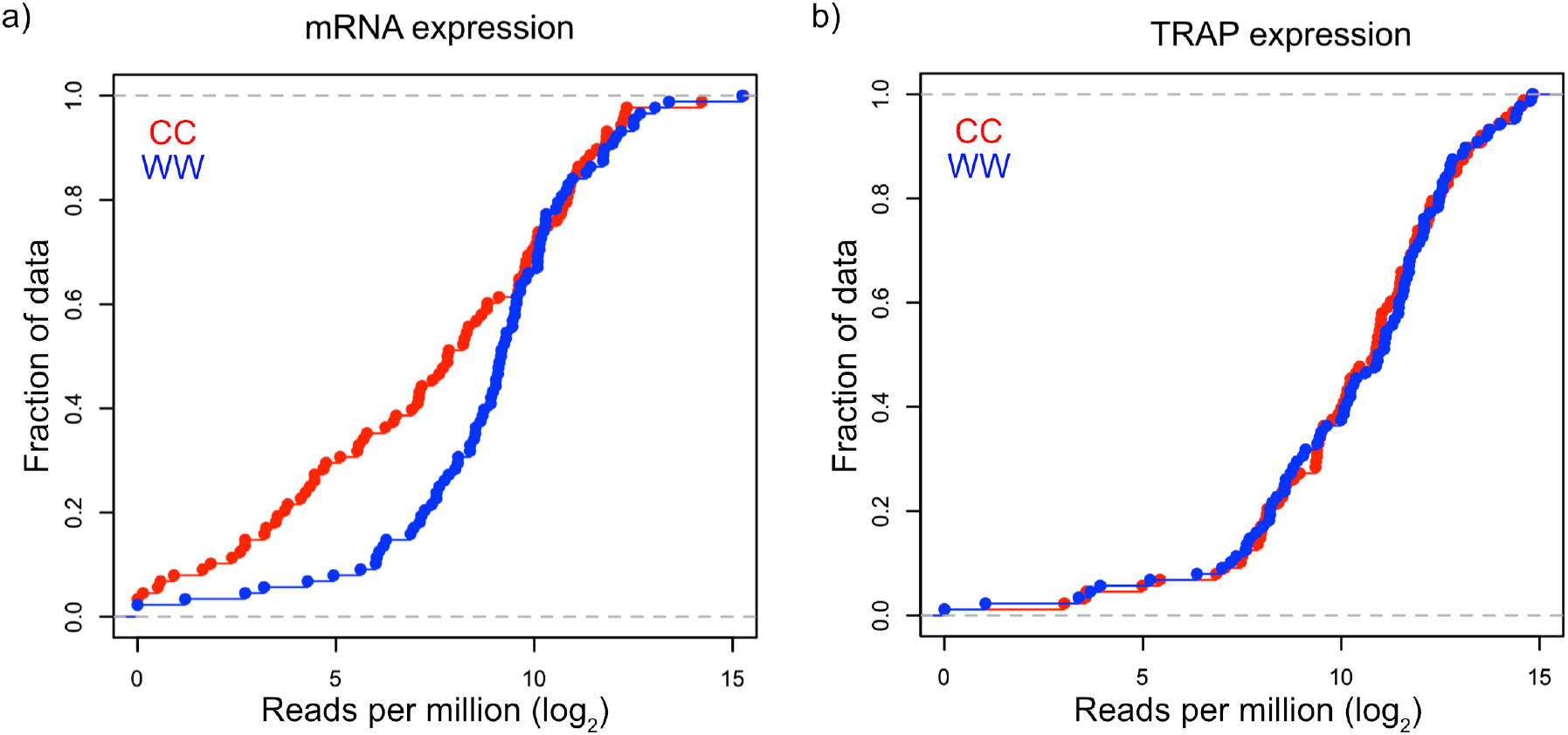
Validation of down-regulation of transcripts and maintained ribosomal loading in Tsc1-KO PCs. a) This cumulative distribution function shows the distribution of total mRNA expression from sorted PCs for 150 FMRP target genes, previously found to be down-regulated in mutant PCs, between mutant (red) and control (blue) PCs. b) This cumulative distribution function shows the TRAP values for the same 150 genes between mutant (red) and control (blue) PCs. Despite the obvious down-regulation of many genes at the total mRNA level, the ribosomal loading of these transcripts appears mostly unchanged.

### SHANK2 expression is decreased in *Tsc1*^c/c^ PCs

Given the competing changes at the transcriptional and ribosomal level, we investigated the resulting protein levels for one of the targets, *Shank2. Shank2* is an ASD candidate gene, a known FMRP target, and has been implicated in proper PC synapse formation (Darnell et al., 2011; Peter et al., 2016; Satterstrom et al., 2020). In addition, it was a top hit in the differential expression analysis but showed no change at the level of ribosomal binding between mutant and control PCs. To characterize the effect of these alterations on the protein level, we performed immunohistochemistry of SHANK2 in the cerebellum of *Tsc1*^w/w^ and *Tsc1*^c/c^ L7Cre;L7GFP animals (N=4 animals of each genotype). We used confocal microscopy to evaluate the intensity of the SHANK2+ punctae in the molecular layer (containing PC dendrites) of mutant and control PCs. Interestingly, we observed a significant decrease in the intensity of SHANK2+ punctae in mutant PCs compared to control (Figure 6). These data suggest that the increase ribosomal binding efficiency of the *Shank2* transcript in *Tsc1*-deficient PCs is unable to overcome its overall down regulation, resulting in a decreased level of the protein at synapses.

**Figure 6:**
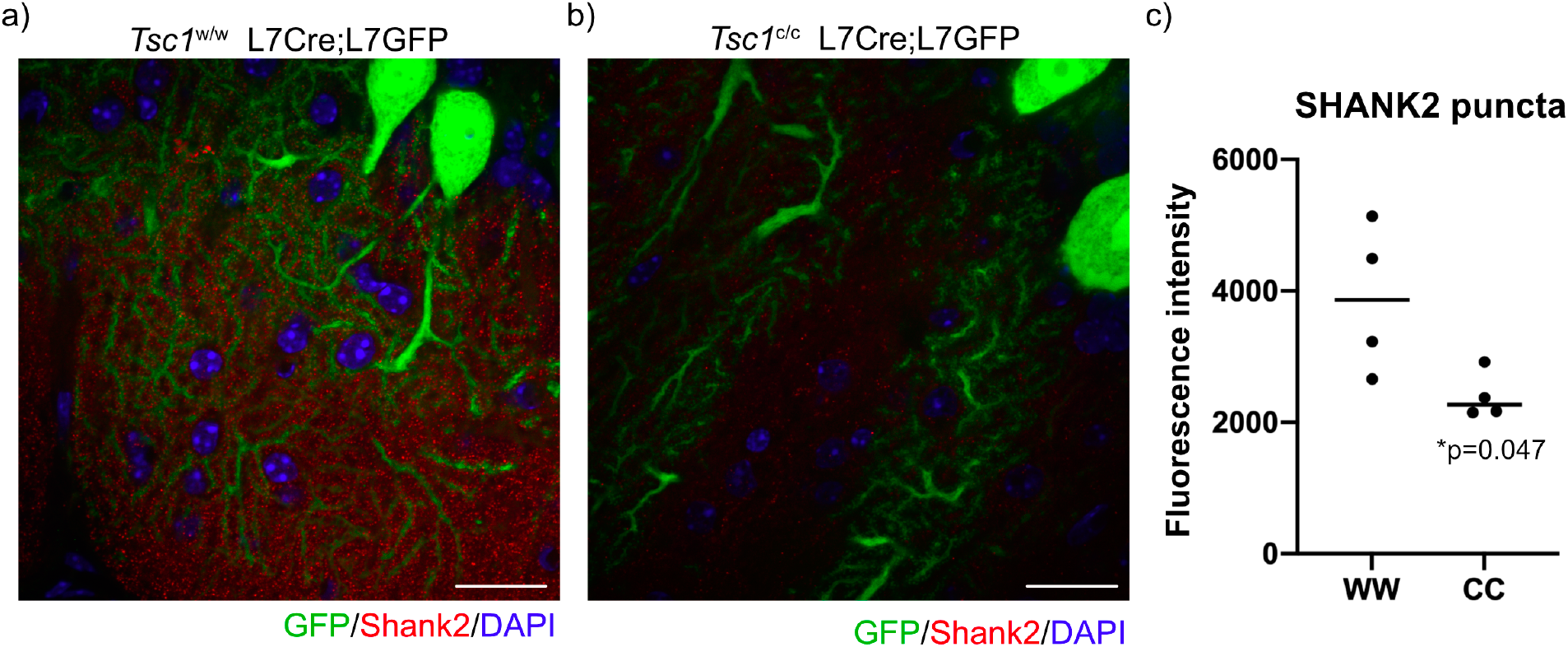
SHANK2 staining in mutant and control PCs. Confocal images of staining of SHANK2 in a) *Tsc1*^ww^ L7Cre+/L7GFP and b) *Tsc1*^c/c^ L7Cre+/L7GFP animals. SHANK2 is present in punctae along PC dendrites in the molecular layer, and GFP-labeled PCs are shown in green. Scale bars = 20um c) Bar plot showing the quantification of the intensity of SHANK2+ punctae in control and mutant animals. Each data point represents the mean value across 10 images from 3-4 independent slices from the same animal. N=4 animals. p=0.047 (T-test).

## Discussion

The cerebellum and PCs, in particular, have been implicated in the development of abnormal social behavior in TSC (Tsai et al., 2012; Tsai et al., 2018). Therefore, we have endeavored to understand the molecular abnormalities present in TSC1-deficient PCs. We observed that the majority of differentially expressed genes between mutant and control PCs were down regulated. We also found that these down-regulated genes in mutant PCs were enriched in FMRP targets and showed reduced coverage at the 3’ region, suggestive of increased RNA decay. In contrast, we did not observe a reduced ribosomal binding of these genes, implicating processes that facilitate more efficient loading of these transcripts onto ribosomes. Finally, we observed that the FMRP target, *Shank2*, showed reduced expression in PC dendrites, implying that compensatory increases in ribosomal binding efficiency may be insufficient to overcome overall reduced transcript levels. These data suggest that dysregulation of FMRP target genes at the transcript level, rather than the ribosomal binding level, may drive molecular dysregulation in *Tsc1* mutant PCs and underlie the development of synaptic abnormalities in these neurons.

We observed that there was a notable bias towards down regulation of genes in mutant PCs, and we found that this was driven by exaggerated reduction of the coverage of the 3’ region, especially the 3’ UTR. Prior studies have reported that mTOR can affect RNA stability, but most of these studies showed that treatment with rapamycin reduced stability of certain transcripts (Banholzer et al., 1997; Hashemolhosseini et al., 1998; Albig and Decker, 2001). Rapamycin reduced the stability of specific transcripts in yeast through increasing the rate of deadenylation, although the mediator of this effect was not identified (Albig and Decker, 2001). Inhibition of mTORC1 with rapamycin was also shown to facilitate RNA degradation through the nonsense mediated decay pathway, but this effect was shown to be associated with alterations in translation machinery associated with the 5’ cap (Martinez-Nunez et al., 2017). It is likely that our observations contradict these earlier studies because none of the prior studies assayed hyperactive mTORC1 or were performed in neurons, and the downstream machinery may be substantially different in mutant PCs. Canonically, RNA degradation is preceded by deadenylation of the transcript, rendering the molecule susceptible to exonuclease activity from either the 5’ or 3’ end (Labno et al., 2016). We hypothesize that 3’->5’ decay activity is increased in mutant PCs because of the reduction in coverage in the 3’ region of transcripts. The best characterized 3’->5’ exonucleases are *Dis3, Dis3L*, and *Dis3L2*, which differ in their association with the RNA exosome complex (Labno et al., 2016; Dos Santos et al., 2018). Interestingly, 20 out of 24 genes associated with the exosome show modest but non-significant increases in expression in *Tsc1*^c/c^ PCs in this dataset (Supplemental table 5). DIS3 was also found to be upregulated due to ER stress in *C. elegans* (Sakaki et al., 2012), and we have found that TSC2-deficient neurons show evidence of ER stress (Di Nardo et al., 2009). In addition, the RNA exosome complex is involved in ribosomal RNA maturation (Kobylecki et al., 2018; Pirouz et al., 2019), and mTORC1 is known to increase ribosomal biogenesis (Gentilella et al., 2015). Therefore, it is possible that mTORC1 may induce the RNA exosome by stimulating the production of ribosomes, and the RNA decay phenomenon that we observe could be a side effect of this process.

Another striking finding of this study is the maintenance of ribosomal binding of transcripts despite down regulation of the gene at the transcript level. These data suggest that the primary abnormality in mutant PCs may be at the transcript level, while the increase in relative ribosomal binding is a compensatory effect. Interestingly, the genes that show this pattern of reduced transcript level with maintained ribosomal binding are highly enriched in targets of FMRP. We also observed a similar decreased expression of FMRP target transcripts in iPSC-derived PCs with mutations in *TSC2* (Sundberg et al., 2018), corroborating the findings of this study. FMRP is an RNA binding protein that is lost in Fragile X Syndrome, which is characterized by high rates of intellectual disability and ASD (Niu et al., 2017). FMRP is known to be a repressor of translation (Feng et al., 1997; Brown et al., 2001), but its effects on the ribosomal binding of specific transcripts are less clear (Thomson et al., 2017). In addition, FMRP can bind to ribosomes and reduce their ability to synthesize new proteins, suggesting a more general effect on translation (Chen et al., 2014). Therefore, it is possible that dysfunction in FMRP could facilitate binding of specific transcripts to ribosome despite their reduced abundance at the total transcript level. FMRP has also been shown to play a role in mRNA stability (Zalfa et al., 2007; De Rubeis and Bagni, 2010), which suggests that it could also contribute to the increased RNA degradation and down-regulation of its target transcripts. A recent study demonstrated that loss of FMRP can preferentially destabilize transcripts in neurons based on their codon optimality (Shu et al., 2020). In contrast, another recent study that examined hippocampal slices from *Tsc2*+/- mice reported that the expression of FMRP targets was increased and their ribosomal binding was decreased (Hien et al., 2020). It is possible, and in fact likely, that there are cell type differences in the functions of TSC1/TSC2 and FMRP in the brain leading to the discrepancy between this prior study and our data. Our study further reinforces the connection between mTOR and FMRP and highlights the role of mTOR hyperactivation in altering both the total abundance and ribosomal binding of FMRP target genes.

Given the compensatory increase in relative ribosomal binding of FMRP targets to control levels, it is unclear whether and how these abnormalities contribute to neuronal dysfunction in *Tsc1-*deficient PCs. One interesting gene that shows down regulation at the transcript level but maintained ribosomal binding is *Shank2. Shank2* is a scaffolding protein that is present at the post-synaptic density and highly expressed in PCs (Eltokhi et al., 2018). One study found that deletion of *Shank2* in PCs altered the levels of glutamate receptors in dendritic spines and led to abnormal social behavior in these animals (Peter et al., 2016). Another study that deleted *Shank2* in PCs found alterations in excitation onto PCs but did not observe abnormal social behavior (Ha et al., 2016). These data suggest that proper regulation of *Shank2* in PCs is critical for normal function of cerebellar circuits. We found that the apparently compensated ribosomal binding of *Shank2* is insufficient to overcome its down-regulation at the transcript level, leading to its down-regulation at the protein level. Therefore, these data suggest that reduced SHANK2 and other synaptic proteins that display similar patterns of dysregulation may contribute to the development of altered cerebellar circuitry and abnormal social behavior.

In this study, we performed profiling of the total transcriptome and ribosomally-bound transcripts in PCs with a deletion of *Tsc1*. We observed changes at the transcript level that suggest alterations in RNA degradation and compensatory changes at the ribosome level. These data indicate the presence of novel mechanisms underlying PC dysfunction in TSC, which may be associated with the development of abnormal social behavior. Further study of these pathways may provide new avenues for therapeutics for ASD in TSC and related neurodevelopmental disorders.

## Materials and Methods

### Mice

All animal procedures were carried out in accordance with the Guide for the Humane Use and Care of Laboratory Animals, and all procedures in this study were approved by the Animal Care and Use Committee of Boston Children’s Hospital. All animals were kept in ARCH animal house facility under 12-hour light/dark cycle with food and water available ad libitum. *Tsc1*^c/c^ mice possess loxP sites flanking exons 17 and 18 of the *Tsc1* gene (stock# 005680) and were obtained from Jackson labs (Kwiatkowski et al., 2002). The Rosa26fsTRAP mouse line (stock# 022367) and the L7Cre mouse line (stock# 010536) were obtained from Jackson labs. The L7GFP line was also obtained from Jackson labs (stock# 004690). The L7GFP mice were on a C57BL/6J background and were bred with *Tsc1*^c/w^ L7Cre animals to generate *Tsc1*^c/w^ L7Cre;L7GFP animals, which were bred together to generate *Tsc1*^c/c^ or *Tsc1*^w/w^ L7Cre;L7GFP animals. The Rosa26fsTRAP mice were on a C57BL/6j background and were bred with *Tsc1*^c/w^ L7Cre animals to generate *Tsc1*^c/w^ L7Cre;Rosa26fsTRAP animals. These animals were bred together to generate either *Tsc1*^c/c^ or *Tsc1*^w/w^ L7Cre;Rosa26fsTRAP animals.

### Fluorescence Activated Cell Sorting (FACS)

For isolation of PCs, we used *Tsc1*^w/w^ or *Tsc1*^c/c^ L7Cre/L7GFP animals. The cerebellum was microdissected from either P21 or P42 animals and chopped into small pieces before incubation with papain digestion buffer (1x HBSS medium containing 40 U/ml Papain (Worthington Biochemical cat # LS003126), DNase I 100 units/ml (Worthington Biochemical cat # LK003172) and 30 mM glucose) for 30-45 min at 37C with 5% CO2. Tissue was dissociated into single cells by trituration with a fire-polished glass pipette. Cell suspensions were washed twice with papain wash buffer (1x HBSS supplemented 30mM glucose, and 10% FBS and 1x penicillin /streptomycin antibiotic cocktail). Cells were then resuspended in FACS buffer (1x HBSS with 1x pen/strep solution and 2% BSA (sigma ca # A9576)) and filtered through 40µM cell strainer (Falcon). Cells were centrifuged 200xg for 3 min and suspended into FACS-buffer (1x HBSS and 1% BSA). Cells were sorted with FACS-Aria II (BD Bioscience) using 100µM nozzle setting. Cells were identified from debris based on the forward and side scatter, and doublets were removed using FSC-H and SSC-W/SSC-H. Cells that were GFP positive and PI negative were collected into FACS-buffer, and between 1-2 million cells were collected per animal. Cells washed once with PBS and total RNA isolated using Qiagen mini Plus kit according to manufacturer instructions (Qiagen). RNA quality and integrity were assessed by analysis in Bioanalyzer. RNA samples with RIN values >8 were used for library preparation according to manufacturer’s instructions (SmartSeqv4 RNASeq kit, Clontech) or manual library preparation for Ampliseq analysis (Ion AmpliSeq Lib Kit Plus). From each mouse, an average of 1,500-1,000,000 GFP positive cells were obtained.

### TRAP

To purify ribosome associated RNAs specifically from Purkinje neurons in the cerebellum from *Tsc1*^c/c^ and *Tsc1*^w/w^ animals, we used TRAP methods as previously described (Heiman et al., 2014; Dougherty, 2017). Briefly, 12μl of Purified Recombinant Biotinylated Protein L (Pierce cat #29997, 1ug/ul) was conjugated to 30μl of Streptavidin MyOne T1 Dynabeads (Thermo Fischer Scientific cat # 65601) at room temperature using gentle end-over-end rotation. After collecting beads on the magnet, the beads were washed five times with PBS containing 3% BSA (Jackson Immunoresearch cat #001-000-162). After collecting beads on the magnet, they were washed once with low KCl wash buffer (10mM HEPES pH7.4, 5mM MgCl_2_, 150mM KCl, and 1% NP-40) and further incubated with 100μg total of mouse anti-GFP antibodies (HtzGFP-19F7 and HtzGFP-19C8, Memorial Sloan Kettering Centre, 50μg of each) for 1 hour at room temperature.

To prepare cerebellar lysates, *Tsc1*^w/w^ or *Tsc1*^c/c^ L7Cre+/Rosa26fsTRAP+ animals were decapitated, and the cerebellum from each animal was homogenized in ice-cold homogenization buffer (10 mM HEPES pH 7.4, 5 mM MgCl_2_, 150 mM KCl, 0.5 mM DTT, 100 μg/ml cycloheximide, Rnasin, Superasin, and 1x protease inhibitors, cOmplete EDTA-free from Roche) using 12 strokes of Teflon-glass homogenizers. To clear the nuclei and debris, samples were centrifuged at 2,000x g for 10 mins at 4C. Polysomes were prepared by adding NP-40 (1%) and DHPC (30mM), incubating on ice for 5 minutes, followed by centrifugation at 20,000x g for 15 mins at 4C. For input RNA, a 60μl sample of the supernatant was removed, added to 190ul of low KCl wash buffer and 750μl of Trizol and stored at −80C for RNA isolation. Remaining supernatant was incubated with GFP-coated Dynabeads overnight at 4C with end-over-end rotation. After overnight incubation, anti-GFP beads were captured on the magnet, washed 4 times with high KCl wash buffer (10 mM HEPES pH 7.4, 5 mM MgCl_2_, 350 mM KCl, 1% NP-40, 0.5 mM DTT, Rnasin, and 100 μg/ml cycloheximide), and resuspended in 250μl of low KCl wash buffer and 750μl of Trizol reagent. RNA was isolated using Trizol reagent protocol. To remove contaminating genomic DNA, DNase I (Qiagen cat #) was used from Qiagen RNAeasy Minelute kit. RNA quality was assessed by Bioanalyzer analysis using Pico chip. TRAP RNA from each animal was used to make Illumina ready libraries at the Molecular Biology Core facility at the Dana Farber Cancer Insitute.

### RNA-Seq

Barcoded and pooled libraries were sequenced on Illumina HiSeq 4000 to generate 75 base paired end reads. Quality of RNA-seq data was assessed initially using FastQC (https://www.bioinformatics.babraham.ac.uk/projects/fastqc/), and Trimmomatic (http://www.usadellab.org/cms/?page=trimmomatic) was used to remove adapters and low quality bases. Reads were mapped to the mouse genome (Ensembl GRCm38) using STAR. For sorted PCs, the transcriptome was assembled with Cufflinks. For the TRAP data, the transcriptome assembly from the sorted PCs was used. Gene and transcript quantification were performed using Cufflinks. These data were imported into R for further analysis.

Differential expression from both datasets was calculated using the LIMMA package. Differentially expressed transcripts were also identified using LIMMA using the transcript specific data from Cufflinks. Thresholds for differential expression were p-value < 0.01 and fold change > 2.

Exon coverage was determined directly from the mapped SAM files for each sample using custom Python scripts. This was then converted into transcript coverage by concatenating exons into transcripts that were identified by Cufflinks. Transcripts were then broken into 100 evenly spaced intervals and average number of reads across an interval were normalized to the total number of reads across the entire transcript. For up and down regulated transcripts, the sequences were obtained using the UCSC Table Browser based on coordinates from Cufflinks. Coding and non-coding (5’UTR and 3’UTR) were extracted from these sequences by searching for the longest open reading frame within the sequence. Coverage was then calculated separately for each portion of the transcript (5’UTR, CDS, 3’UTR), and the difference between wildtype and mutant cells was calculated. Gene ontology was performed using DAVID (https://david.ncifcrf.gov/), and non-redundant categories from GO_FAT Cellular Compartment, Biological Process, and Molecular Function were selected.

TE was calculated as described in Thoreen et al., 2012. Briefly, counts from RNA and TRAP were transformed using log_2_(counts + 1), and ΔTE = (TRAP *Tsc1*^c/c^ – TRAP*Tsc1*^w/w^) – (RNA *Tsc1*^c/c^ – RNA*Tsc1*^w/w^). Variance of the ΔTE measure was estimated by recalculating ΔTE one hundred times per gene, while leaving one value out for each dataset. This distribution was summarized using a Z-score for each gene.

### Immunohistochemistry

*Tsc1*^w/w^ or *Tsc1*^c/c^ L7Cre/L7GFP (N=4 for each genotype) were perfused at P42 with saline and then 4% PFA. Brains were dissected, post-fixed in 4% PFA for 24 hours, and then placed in 30% sucrose for at least 24 hours. Floating sections (40um) were cut on a cryostat and stained with anti-GFP (Thermo Fisher A10262) and anti-SHANK2 (Thermo Fisher PA5-78652). Images were obtained on a spinning disk confocal microscope with 10 images per animal. SHANK2 punctae were identified and quantified using custom ImageJ macros.

### Statistical methods

Count data from RNA sequencing and TRAP were imported into R for further analysis. Differential expression from both datasets was calculated using the LIMMA package, and the thresholds for differential expression are indicated in the text. Differences in coverage between sections of transcripts from up and down-regulated genes was performed using an ANOVA with Tukey’s post-hoc test. Other statistical tests were performed in R and are indicated in the text. For IHC data, the average SHANK2 punctae intensity was imported into PRISM, and the comparison between wildtype and mutant was performed using a T-test.

## Acknowledgements

We would like to thank Elizabeth Bainbridge and Sarika Gurnani for help with mouse breeding. The Sahin lab has received grant funding from the U.S. Army Medical Research Tuberous Sclerosis Complex Research Program (W81XWH-15-1-0189). Boston Children’s Hospital Intellectual and Developmental Disabilities Research Center (BCH IDDRC, U54HD090255). JD was supported by Roche Postdoctoral Fellowship (RPF) Program. KDW is funded by the Neuroscience Research Training Scholarship from the American Academy of Neurology and NIH (5K08NS112598-02).

---

**Supplemental Figure 1:**
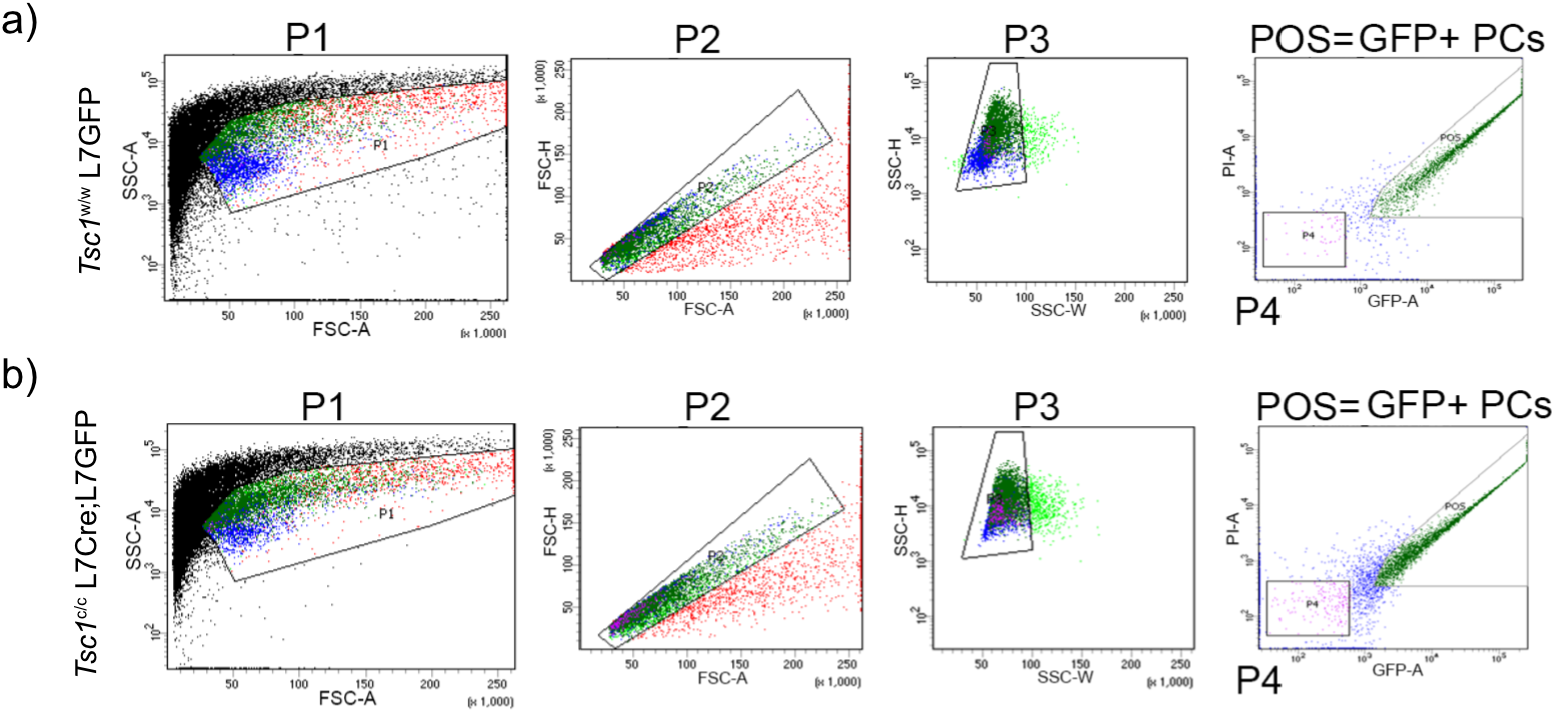
Gating strategy for FACS isolation of GFP labeled PCs. The gating for isolation of GFP-positive PCs is shown for a) Tsc1w/w L7GFP and b) Tsc1c/c L7Cre;L7GFP animals. Cells were identified from debris based on the forward and side scatter (P1). Single cells were identified using FSC-H (P2) and SSC-H/SSC-W (P3). Finally, GFP-positive/ PI-negative were cells were selected and sorted into separate tubes (POS).

